# Longitudinal fundus imaging and its genome-wide association analysis provide evidence for a human retinal aging clock

**DOI:** 10.1101/2022.07.25.501485

**Authors:** Sara Ahadi, Kenneth A. Wilson, Boris Babenko, Cory Y. McLean, Drew Bryant, Orion Pritchard, Enrique M. Carrera, Ricardo Lamy, Jay M. Stewart, Avinash Varadarajan, Marc Berndl, Pankaj Kapahi, Ali Bashir

## Abstract

Biological age, distinct from an individual’s chronological age, has been studied extensively through predictive aging clocks. However, these clocks have limited accuracy in short time-scales. Deep learning approaches on imaging datasets of the eye have proven powerful for a variety of quantitative phenotype inference tasks and provide an opportunity to explore organismal aging and tissue health.

Here we trained deep learning models on fundus images from the EyePACS dataset to predict individuals’ chronological age. These predictions led to the concept of a retinal aging clock, “eyeAge”, which we employed for a series of downstream longitudinal analyses. eyeAge was used to predict chronological age on timescales under a year using longitudinal fundus imaging data from a subset of patients. To further validate the model, it was applied to a separate cohort from the UK Biobank. The difference between individuals’ eyeAge and their chronological age, hereafter “eyeAgeAccel”, was computed and used for genome-wide association analysis (GWAS).

EyeAge predicted chronological age more accurately than other aging clocks (mean absolute error of 2.86 and 3.30 years on quality-filtered data from EyePACS and UKBiobank, respectively). Additionally, eyeAgeAccel was highly independent of blood marker-based measures of biological age (e.g. “phenotypic age”), maintaining an all-cause mortality hazard ratio of 1.026 even in the presence of phenotypic age. Longitudinal studies showed that the resulting models were able to predict individuals’ aging, in time-scales less than a year, with 71% accuracy. The individual-specific component to this prediction was confirmed with the identification of multiple GWAS hits in the independent UK Biobank cohort. The knockdown of the fly homolog to the top hit, *ALKAL2*, which was previously shown to extend lifespan in flies, also slowed age-related decline in vision in flies.

In conclusion, predicted age from retinal images can be used as a biomarker of biological aging that is independent from assessment based on blood markers. This study demonstrates the potential utility of a retinal aging clock for studying aging and age-related diseases and quantitatively measuring aging on very short time-scales, opening avenues for quick and actionable evaluation of gero-protective therapeutics.

## Introduction

Aging causes molecular and physiological changes throughout all tissues of the body, enhancing the risk of several diseases.^1^ Identifying specific markers of aging is a critical area of research, as each individual ages uniquely depending on both genetic and environmental factors.^2^ While a variety of aging clocks have recently been developed to track the aging process, including phenotypic age^3^ (blood markers based on an individual’s chronological age) and an epigenetic clock derived from DNA methylation,^4^ many require in-depth molecular analysis and potentially invasive cell or tissue extraction.

A growing body of evidence suggests that the microvasculature in the retina might be a reliable indicator of the overall health of the body’s circulatory system and the brain. Changes in the eyes accompany aging and many age-related diseases such as age-related macular degeneration (AMD),^5^ diabetic retinopathy,^6^ and neurodegenerative disorders like Parkinson’s^5,7^ and Alzheimer’s.^8^ Eyes are also ideal windows for early detection of systemic diseases by ophthalmologists, including AIDS,^9,10^ chronic hypertension,^11^ and tumors.^12^ This broad utility is perhaps unsurprising, as any subtle changes in the vascular system first appear in the smallest blood vessels, and retinal capillaries are amongst the smallest in the body.

The subtle changes induced in these small vessels often go undetected by even the most sophisticated instruments, necessitating the use of better approaches involving deep learning. Fundus imaging has proven to be a powerful and non-invasive means for identifying specific markers of eye-related health. Deep-learning was initially employed to predict diabetic retinopathy from retinal images at accuracies matching, or even exceeding, experts.^13^ Since then, retinal images have been employed to identify at least 39 fundus diseases including glaucoma, diabetic retinopathy, age-related macular degeneration (AMD),^11,14^ cardiovascular risk,^15^ chronic kidney disease,^16^ and, most recently, in predicting age.^17^ Given its non-invasive, low-cost nature, retinal imaging provides an intriguing opportunity for longitudinal patient analysis to assess the rate of aging.

Here we use deep learning models to predict chronological age from fundus retinal images, hereafter “eyeAge”, and use the deviation of this value from chronological age, hereafter eyeAgeAccel, for mortality and association analyses. We train this model on the well-studied EyePACS datasets and apply it on both the EyePACS and UKBiobank cohorts. Together, our results suggest that the trajectory of an individual’s biological age can be predicted in timelines under a year and that statistically significant genome-wide associations are possible. Enrichment analysis of top GWAS hits as well as experimental validation of the *Drosophila* homolog of *ALKAL2*, a gene in the top GWAS locus, indicates genetic markers of visual decline with age and demonstrates the potential predictive power of a retinal aging clock in assessing biological age.

## Results

### Prediction of age from fundus images

Figure 1 summarizes the analysis workflow for the study. Using the EyePACS dataset, we trained a fundus image model on 217,289 examples from 100,692 patients and tuned it on 54,292 images from 25,238 patients. These models were employed for longitudinal analysis of repeat patients and also applied on the UK Biobank dataset (119,532 images) which had a notably distinct demographic distribution (Table 1). In both analyses, we took the average of the predictions between the left and right eye from a single visit to infer age (See Methods).

**Table 1.** Characteristics of patients in the development and validation sets (before filtering).

|  | Development set (EyePACS) |  | Test set (UK Biobank) |
| --- | --- | --- | --- |
|  | Train | Tune |  |
| Patients | 100,692 | 25,238 | 64,019 |
| Images | 217,289 | 54,292 | 119,532 |
| Ethnicity | Black: 11908 [7%] | Black: 3040 [7%] | Black: 1540 [1%] |
|  | Asia Pacific Islander: 11842 [7%]<br>White: 22539 [13%]<br>Hispanic: 125595 [71%]<br>Native American: 1791 [1%]<br>Other: 3809 [2%] | Asia Pacific Islander: 2923 [7%]<br>White: 5657 [13%]<br>Hispanic: 31521 [71%]<br>Native American: 426 [1%]<br>Other: 918 [2%] | Asia Pacific Islander: 4183 [4%]<br>White: 107967 [91%]<br>Hispanic: 0 [0%]<br>Native_american: 0 [0%]<br>Other: 5015 [4%] |
| Gender | Female: 127075 [59%]<br>Male: 90128 [41%] | Female: 31743 [58%]<br>Male: 22531 [42%] | Female: 65739 [55%]<br>Male: 53793 [45%] |
| Age | median=55.13<br>mean=54.21<br>std=11.50 | median=55.19<br>mean=54.20<br>std=11.46 | median=57.94<br>mean=56.85<br>std=8.18 |

**Figure 1.**
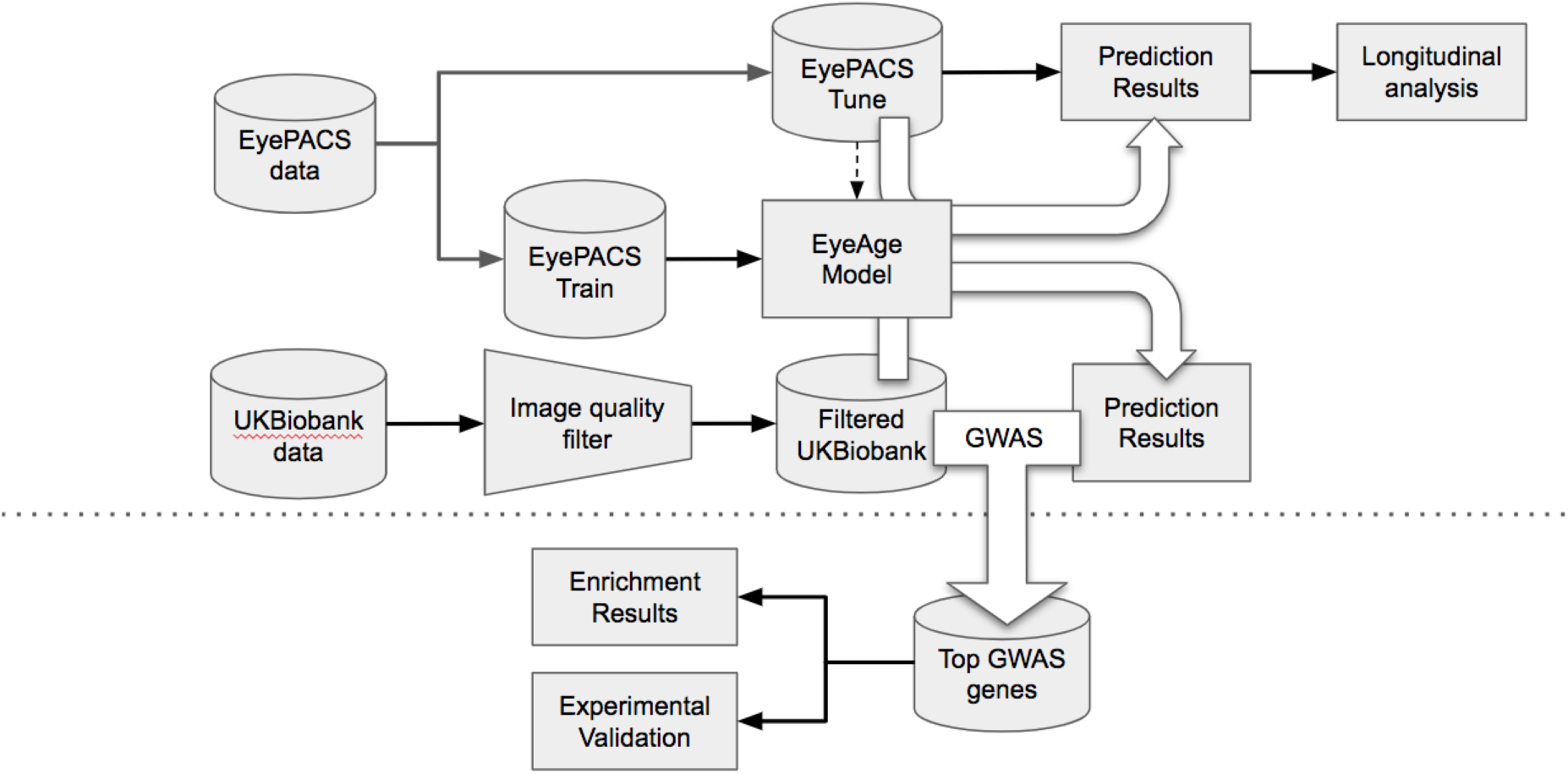
Schematic of analysis pipeline. EyePACS images were split into train and tune sets based on the patient. The model was then trained with the final model step being selected via the tune set. Prediction results on the EyePACS tune set were used for longitudinal analysis of aging. After filtering for image quality, inference was performed with the same model on the UKbiobank dataset and filtering for image quality, and the resulting eyeAgeAccel was used for GWAS analysis. Enrichment analysis was performed on the GWAS hits with a homolog of the top gene (*ALKAL2*) validated experimentally in *Drosophila*.

The model showed a strong correlation between chronological age and eyeAge (eyeAge) in both the EyePACS (0.95) and UK Biobank (0.87) datasets (Supplementary Figure 1). Using mean absolute error (MAE) to assess the fidelity of the aging clock showed that the model performed favorably on both datasets (2.86 and 3.30, respectively, after quality filtering) relative to previous studies.^17–20^ Next, we evaluated the efficacy of our predictions in one to two year time scales using longitudinal datas. Using the EyePACS Tune dataset, we restricted ourselves to data from patients with exactly two visits (1,719 subjects) and examined the models’ ability to order the two visits over multiple time scales. Note that no longitudinal information about patients was specifically used to train or tune the model to predict chronological age. Figure 2A shows that the model correctly ordered 71% of visits within a year with an MAE less than 2 years. In both metrics the fidelity decreased in older groups and with smaller age gaps.

**Figure 2.**
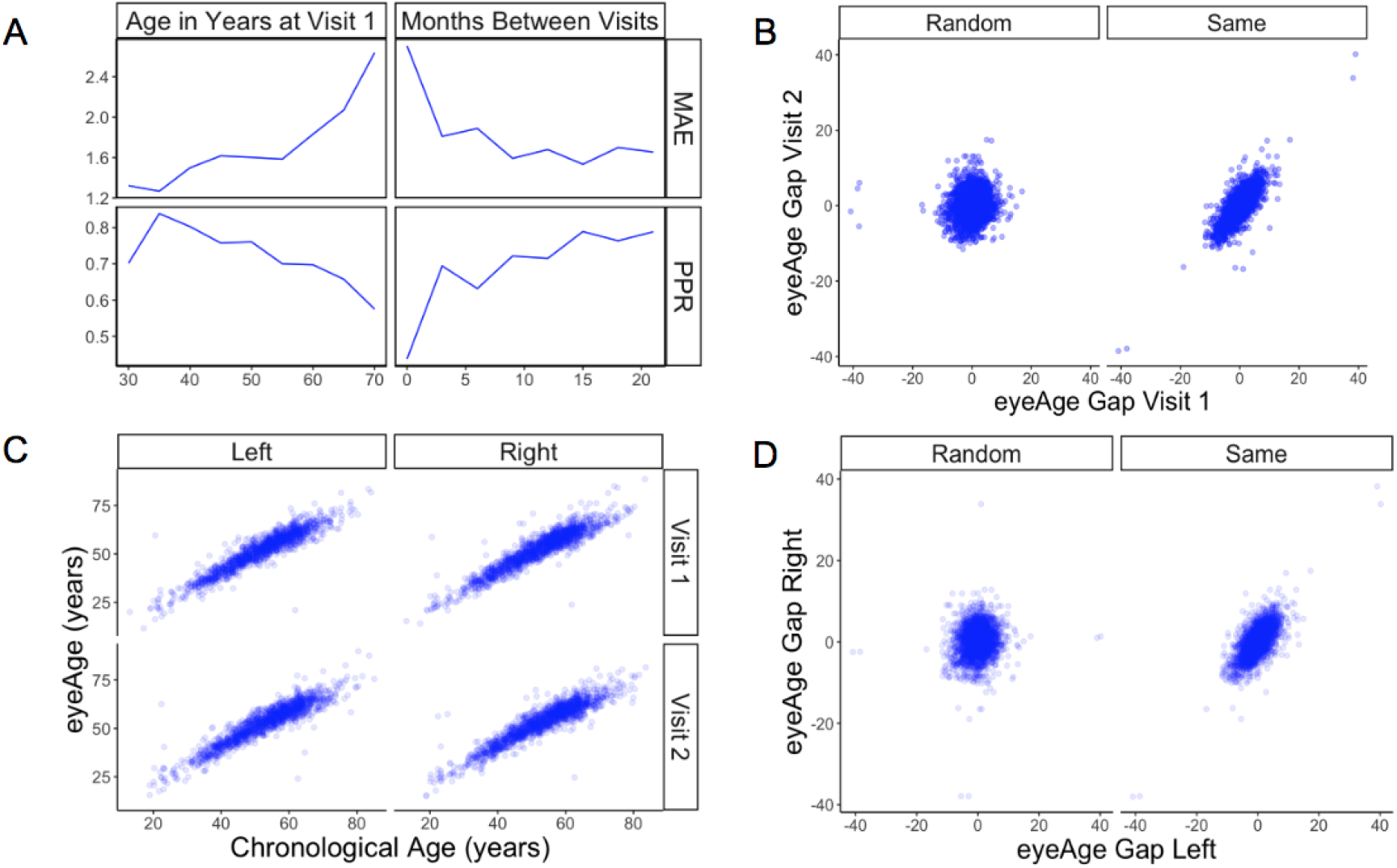
Longitudinal analysis of patients with exactly two visits. (A) Changes of PPR (positive prediction ratio: the ratio of data whose eyeAge increased between subsequent visits) and MAE (mean absolute error) calculated on the same individual in relationship to chronological age at the first visit (left) and time between longitudinal visits (right). (B) Scatter plots representing correlation between eyeAge Gap (difference between predicted age and chronological age) of two consecutive visits from an individual (Same) or two consecutive visits from two different individuals (Random). (C) Correlation of eyeAge and chronological age between left and right and two consecutive visits of the same individual. D) Scatter plots representing the correlation of left and right eyeAge Gap from the same or two random individuals

To understand if this effect was simply a result of the noise of our innate age prediction, we performed an age-matched control experiment. We compared correlations between data points of one individual to data from a random pair of age-matched individuals (see Methods). Comparisons were performed between each eye and timepoint. For all comparisons, the robust correlation observed within an individual’s data was lost in data between time-matched individuals (Figure 2B,D). Additionally, the positive predictive percentage and MAE exhibited reduced performance, 55% and 3.6 years (Supplemental Figure 2), suggesting a reproducible, individual-specific eyeAge component. To further explore this individual-specific component, Figure 2C compares eyeAge and chronological age within an individual between eyes and timepoints, showing strong correlation in each quadrant.

### Testing the model in UK Biobank cohort

We next applied our EyePACS-trained eyeAge model to the UKBiobank dataset. The UKBiobank cohort included retinal fundus images from 64,019 patients as well as extensive clinical labs and genomic data. These clinical markers enabled comparison of eyeAge with “phenoAge”, a clinical blood marker-based aging clock.^3^ The observed 0.87 correlation between eyeAge and chronological age in the UK Biobank cohort was consistent with (and slightly higher than) the observed correlation of phenoAge and chronological age (0.82) (Figure 3A). Notably, the correlation between phenoAge and eyeAge was substantially lower (0.72) and, in fact, roughly equivalent to the product of their respective correlations with chronological age, suggesting that they were largely independent (Figure 3B). To further explore this relationship, we performed a Cox proportional hazards regression analysis (via the lifelines package, https://github.com/CamDavidsonPilon/lifelines) to assess mortality risk.^21^The hazard ratio for eyeAgeAccel was was statistically significant when adjusting for gender (1.09, p-value=1.6e-53), gender and age (1.04, p-value=1.8e-4), and gender, age, and phenoAge (1.03, p-value=2.8e-3) (Figure 3D).

**Figure 3.**
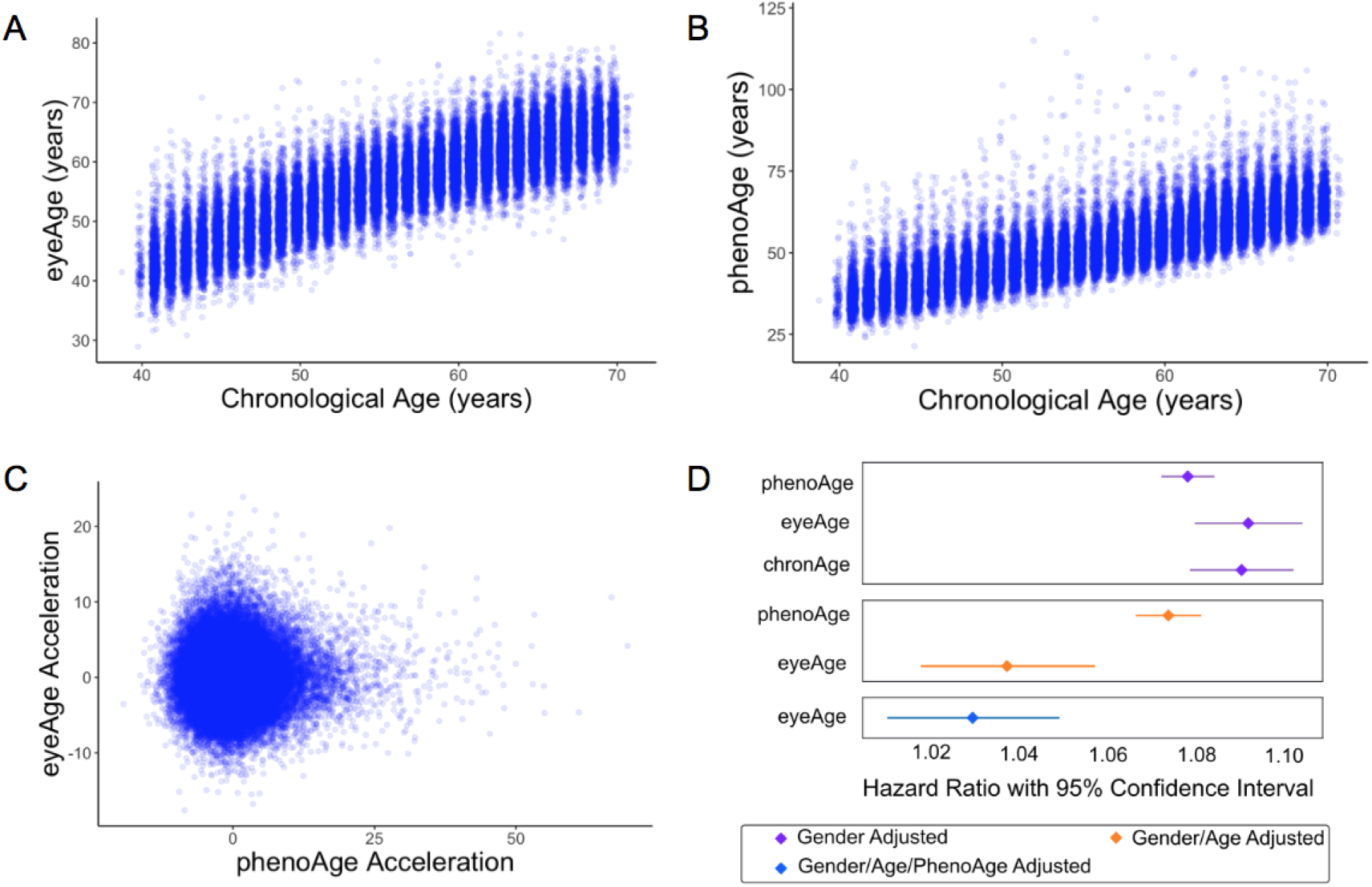
Relationships between eyeAge, phenoAge, and chronological age. (A) Correlation between eyeAge and chronological age (Pearson ρ = 0.86). (B) Correlation between phenoAge and chronological age (Pearson ρ =0.82). (C) Correlation between eyeAgeAcceleration and phenoAgeAcceleration (Pearson Rho = 0.12). (D) Forest plot of all-cause mortality hazard ratios (diamonds) and confidence intervals (lines) for the UKBiobank dataset. Purple lines are adjusted only for gender; orange lines are adjusted for gender and age; blue lines are adjusted for gender, age, and phenoAge.

### GWAS and experimental validation of ALK

Based on these patient-specific effects and independence from phenoAge, a GWAS was conducted to identify genetic factors associated with eyeAge. Prior to performing the GWAS we needed a value that represented a property intrinsic to the patient, and thus more independent of an individual’s age at the time of collection than eyeAge. Therefore, we computed the residuals from linear models that regressed both phenoAge and eyeAge to chronological age, as described previously,^3^ yielding phenoAge acceleration (phenoAgeAccel) and eyeAge acceleration (eyeAgeAccel). We subsetted the cohort to individuals of European ancestry, performed genotype quality control, and utilized a single eyeAgeAccel value per individual, resulting in a cohort of 45,444 individuals for GWAS analysis. GWAS was performed using BOLT-LMM (see Methods) with chronological age, sex, genotyping array type, the top five principal components of genetic ancestry, and indicator variables for the six assessment centers used for the imaging as covariates. Full GWAS summary statistics are available in Supplemental Table 1.

Genomic inflation was low (1.05) (Supplemental Figure 4). The stratified linkage disequilibrium (LD) score regression-based intercept was 1.02 (SEM=0.01), indicating that polygenicity, rather than population structure, drove the test statistic inflation. The SNP-based heritability was 0.11 (SEM=0.02), an appreciable fraction of the estimated broad-sense heritability of biological age (27-57% via twin and family studies). The GWAS identified 38 independent suggestive hits (R^2^ ≤ 0.1, p ≤ 1×10^−6^) at 28 independent loci, 12 of which reached genome-wide significance (p ≤ 5×10^−8^) (Figure 4, Supplemental Table 2).

**Figure 4.**
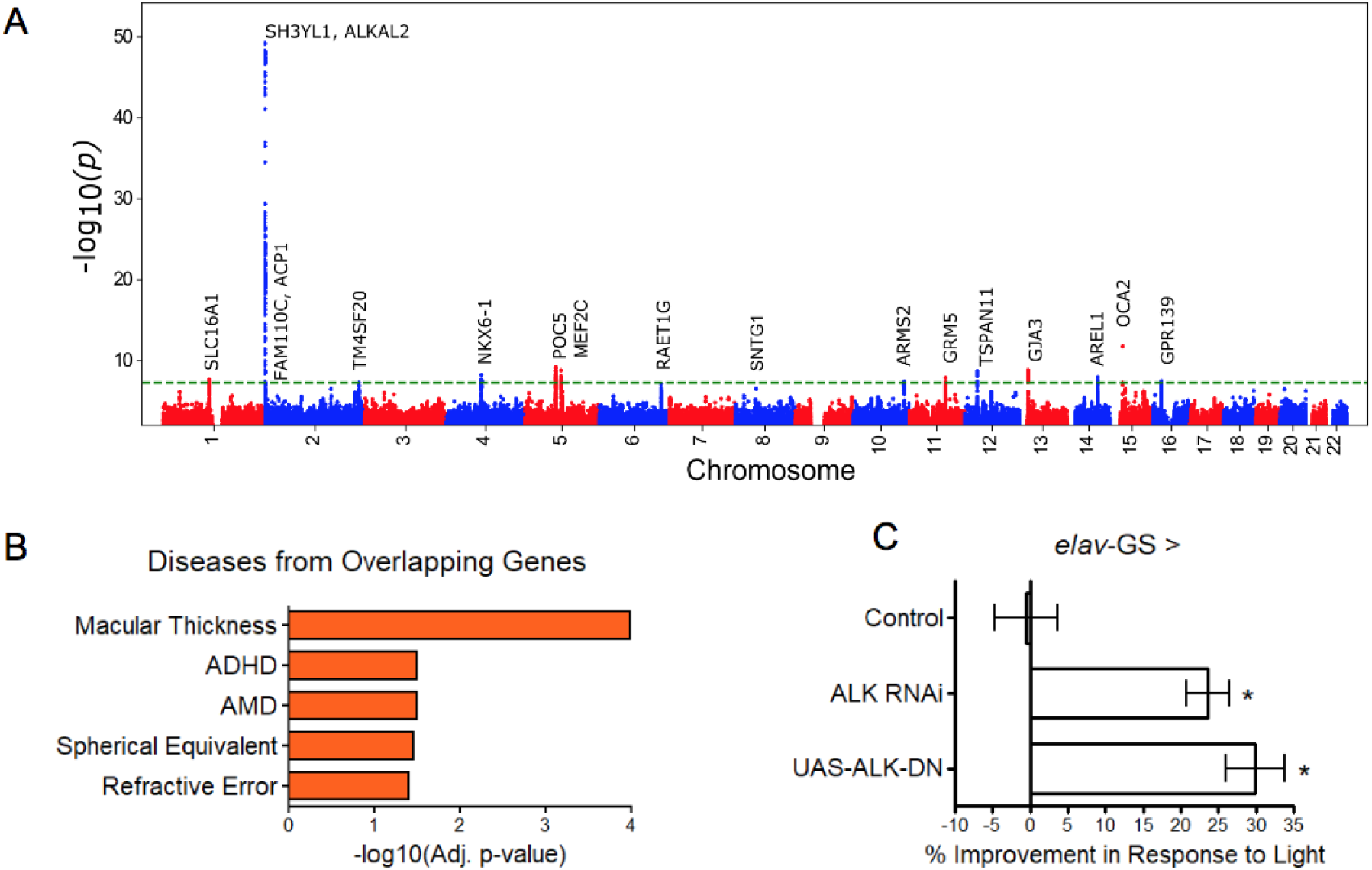
GWAS analyses and experimental validation. (A) Manhattan plot representing significant genes associated with eyeAgeAcceleration. (B) P-values for enriched pathways: Macular thickness, ADHD (attention deficit hyperactivity disorder), AMD (age-related macular degeneration), spherical equivalent, and refractive error. (C) Assessment of visual performance of transgenic and control flies with age. P-value is relative to control (*= p < 0.05).

Many of the hits were associated with eye function and age-related disease (truncated list of candidate hits summarized in Supplemental Table 4). The most significant locus spanned 650 kb and included three genes in a highly significant LD block: *SH3YL1, ACP1*, and *ALKAL2* (Supplemental Figure 5). The *SH3YL1* gene has recently been implicated as a biomarker for nephropathy in type 2 diabetes,^22^ whereas *ALKAL2* enables protein tyrosine kinase activity.^23^ In other significant gene candidates, we identified variants in the genes *OCA2, POC5*, and *GJA3*, which have all been implicated in eye development and function. *OCA2* specifically is known to be important for eye pigmentation,^24^ whereas *POC5* is linked to AMD.^25^ *GJA3* has been implicated in age-related cataract development.^26^ *MEF2C* has reported roles in numerous age-related conditions, including Alzheimer’s disease^27^ and muscle wasting in cancer^28^ and *GRM* is associated with age-related hearing loss.^29^ Additional candidates are reported to be involved in cancer prognosis and progression, including *TSPAN11*,^30^ *NKX6-1*,^31^ and *SLC16A1*.^32^

Gene enrichment analysis^33^ identified significant associations (adjusted p < 0.05) between our gene candidates and macular thickness and degeneration, as seen in previous human GWAS studies^34^ and cataract formation (Elsevier pathway collection),^35^ as well as non-eye related diseases such as bone mineralization, tumor suppression, and Amyloid Precursor Protein pathways (Biocarta).^36^ Gene Ontology (GO) term analysis of our gene candidates revealed significant enrichment (adjusted p < 0.05) for protein tyrosine kinase activator activity, gap junction channel activity, and wide pore channel activity (Figure 4B).

Sum of single effects regression^37^ was used to identify putative causal variants for each locus (Supplemental Table 3). In the most significant locus (Supplemental Figure 5), we identified the deletion variant rs56350804 as the single variant with a posterior inclusion probability (PIP) above 0.45 (rs56350804 PIP=0.9998). While rs56350804 is intronic to *SH3YL1*, expression quantitative trait locus (eQTL) analysis by the Genotype-Tissue Expression consortium identified significant eQTL between rs56350804 and each of *SH3YL1, ACP1*, and *ALKAL2* (GTEx Consortium 2020). In particular, the *ALKAL2* gene had its expression modulated by rs56350804 in cervical spinal cord tissue (p=3.0×10^−16^), and inhibition of the *Drosophila* homolog of *ALKAL2, Alk*, has been shown to extend lifespan,^23^ making it a good candidate for exploring a potential role in visual function.

Previously, *D. melanogaster* has been used to study the impact of aging interventions on retinal health by using the phototaxis index, a fly’s ability to be attracted toward light.^38^ We used *D. melanogaster* to observe visual decline via phototaxis with transgenic *ALK* inhibition. We crossed the pan-neuronal RU486-inducible Gal4 driver *elav*-*Gal4*-GS with UAS-*Alk*^RNAi^ flies or UAS-*Alk*^DN^ to determine the effects of neuron-specific *Alk* inhibition. Both transgenic interventions resulted in significantly increased visual performance with age, whereas background controls showed no change in performance with RU486 treatment (Figure 4C). These results support the implication from the GWAS that *ALK* influences the aging of the visual system.

## Discussion

Retinal health has long been an important factor for visual aging, manifested as glaucoma, AMD, and other age-related retinal diseases, but until recently it was not known whether it could be indicative of overall health and aging. In this study, we applied deep learning models for predicting an individual’s age from retinal fundus images and showed that these predictions may be informative for tracking aging patterns longitudinally. While other cellular and blood-related molecular markers of aging have recently been identified, these are at times invasive and, although accurate, take a long time to develop.^20^ Other aging clocks from blood,^20,39^ saliva,^40^ skin,^40,41^ muscle,^42^ and liver^43^ showed an MAE deviating 4-8 years from the actual age. More dynamic markers such as proteins and metabolites can track aging in shorter time intervals but are still limited to 2-4 years.^2,43,44^ In contrast, using deep learning models on retina fundus images, we were able to predict changes in aging at a granularity of less than a year. These small time-scales, and relative low-cost of imaging, makes eyeAge promising for longitudinal studies.

Correlation and hazard ratio analyses from our study suggest that eyeAge and phenotypic age are conditionally independent given chronological age. Therefore, eyeAge is a potential biomarker that reflects a layer of biological aging not included in blood markers. This is supported by our GWAS findings; different genes were associated with eyeAgeAccel compared to phenoAgeAccel. Similar to other aging clocks (such as DNA-methylome), eyeAge underperforms phenotypic age in mortality prediction. This is likely because the biomarkers used to calculate phenotypic age were explicitly selected based on their ability to predict mortality. New algorithms that incorporate blood markers and retinal clocks have the potential to be better predictors of morbidity and mortality.

Our GWAS identified candidate genes associated with several eye- and age-related functions, such as *POC5*^*25*^ and *GJA3*.^26^ Additional significant candidates had previously identified functions that are not restricted to the eye but are still related to age, e.g. *MEF2C* being associated with Alzheimer’s disease ^27^ and multiple candidates (TSPAN11, NKX6-1, SLC16A1, RAET1G, SNTG1, ARRDC3, RASSF3, DIRC3, and GCNT3) associated with cancer (Supplemental Table 4). These suggest that eyeAge may identify general signatures of aging rather than purely eye-related traits. Pathway analyses similarly were split between eye-related pathways and others that were not eye-specific. While we suspect many of the eye-related pathways to have an aging component, some pathways may be enriched artifactually. For example, though melanin biosynthesis has been associated with protection from photodamage,^38^ the predicted quality of fundus images has also been shown to be influenced by eye color.^45^ Notably, an independent group separately identified our top GWAS candidate locus as the most significant locus.^46^ This combined with previous studies showing *ALK* to be important for lifespan extension in flies^23^ and our own experimental validation confirming improved ocular health in a fly homolog, *Alk*, is compelling evidence of a true biological signal in the GWAS.

Taken together, our work reinforces the utility of fundus imaging for evaluating overall health and opens up new opportunities for using it to predict longevity. eyeAge has substantial applications in aging and aging-related diseases, from biomarker application to tracking therapeutics. In particular, the retinal aging clock because of its ease of use, low cost, and non-invasive sample collection, has the unique potential to additionally assess lifestyle and environmental factors implicated in aging. Retinal aging clocks can be immensely valuable to future clinical trials of rejuvenation/anti-aging therapies and for personalized medicine to measure improvements in aging over short periods, not only improving actionability but also enabling rapid iteration.

## Methods

### EyeAge model development

Model development was done on the EyePACS train dataset (Table 1). A deep learning model with an Inception-v3 architecture ^47,48^ was trained to take a color fundus photo as input and predict the chronological age (referred to as chronologicalAge below) using L1 loss. Age values were normalized to have zero mean and unit variance before training (and during inference this normalization is reversed to get back to year units). Model training was stopped after 363,200 steps by looking at performance on the EyePACS tune dataset. The hyperparameters of the model were as follows: the initial learning rate was 0.0001, which was warmed up to 0.001 over 40751 steps; after the warm up phase, the learning rate was decayed by a factor of 0.99 every 13584 steps; dropout was applied to the prelogits at a rate of 0.8; a weight decay of 4e-5 was used. The model backbone was pre-trained using the ImageNet dataset.^47^ As some of the color fundus images in the UKBiobank dataset were of very low quality, we also trained a separate deep learning model to predict image quality, similar to what was reported in our prior work.^49,50^

### EyeAge model evaluation

The model described above was applied to images to predict chronological age. The image quality model described above was used to discard low quality images – reducing the initial 85649 patient (174,057 image) dataset to 66,536 patients (120,368 images). Finally, we restricted the data to the first assessment visit to UKBiobank. This was done to reduce bias associated with image quality differences, as we observed quality differences between images captured in the later follow-up visits. Since these follow-up visits happened several years after the initial assessment, the time to event or censorship is much smaller, and a model could exploit this association. For participants that had images of both eyes passing the quality filter, we averaged the predictions across the two eyes. After these processing steps, we ended up with 55,269 data points total, one per remaining participant. Next, using the predicted eyeAge and the chronologicalAge of the participant at the time of imaging, an “eyeAgeAcceleration” score was calculated for each participant as the residuals of the ordinary least squares regression model “chronologicalAge ∼ eyeAge”.^3^ In order to compare with another well known biological marker of age, phenoAge^3^ was also computed using the values of blood markers available for the participants. PhenoAgeAcceleration was then computed in an analogous manner to eyeAgeAcceleration.

### Method on selection of random set

Figure 2 required identification of matched, random individuals to assess the potential person-specific component of eyeAge predictions. For Supplemental Figure 2, we created matched sets of visit pairs for each patient’s first visit by identifying a randomly matching patient visit that was 0-2 years after a patient’s first visit. To eliminate artifacts due to sampling differences between first and second visits, once we identified a patient’s first visit to match, we constrained its set of potential randomly matched patient visits to only be from second visits. For the longitudinal analysis in 2B (right), individuals were split both by age and by time between visits (using 2 month buckets) and, again, randomly matched. For Figure 2D, the individuals were split evenly in 2 year buckets. Individuals within the same bucket had their left and right predictions compared to one another.

### GWAS

The eyeAgeAccel value defined above was used as the target for GWAS analysis. GWAS analysis was performed on the fundus-based phenotype as described previously.^51^ Briefly, samples were restricted to individuals of European ancestry to avoid confounding effects due to population structure. European genetic ancestry was defined by computing the medioid of the 15-dimensional space of the top genetic principal components in individuals who self-identified as “British” ancestry and defining all individuals within a distance of 40 from the medioid as “European” (corresponding to the 99th percentile of distances of all individuals who self-identified as British or Irish). Samples were further restricted to those who also passed sample quality control measures computed by UK Biobank, i.e. those not flagged as outliers for heterozygosity or missingness, possessing a putative sex chromosome aneuploidy, or whose self-reported and genetically-inferred sex were discordant.

BOLT-LMM v2.3.4 was used to examine associations between genotype and eyeAgeAcceleration in European individuals in the UK Biobank (n=45,444). All genotyped variants with minor allele frequency > 0.001 were used to perform model fitting and heritability estimation. GWAS was performed in genotyped variants and imputed variants on autosomal chromosomes, with imputed variants filtered to exclude those with minor allele frequency (MAF) < 0.001, imputation INFO score < 0.8, or Hardy-Weinberg equilibrium (HWE) P < 1×10^−10^ in Europeans. In total, 13,297,147 variants passed all quality control measures. Covariates included in the association study were chronological age, sex, genotyping array type, the top five principal components of genetic ancestry, and indicator variables for the six assessment centers used for the imaging.

Genome-wide suggestive (p ≤ 1 × 10^−6^) lead SNPs, independent at R^2^≤0.1, were identified using the `−clump` command in PLINK version v1.90b4. The LD reference panel contained 10,000 unrelated UK Biobank subjects of European ancestry (as defined above). To identify distinct non-overlapping loci of association, all variants with R^2^ ≥ 0.1 with a lead SNP were grouped into a “cluster” with that lead SNP, and subsequently clusters within 250 kilobases of each other were merged, with the lowest p-value lead SNP retained as the locus representative. Putative causal variants were identified using susieR version 0.9.0. At each locus containing at least 10 variants in LD, the `susieR::susie_suff_stat` function was used to estimate posterior inclusion probabilities for each variant in the locus, using the same LD reference panel as was used to generate loci and with a maximum of L=10 causal variants per locus and 200 iterations of coordinate ascent.

### Validation of Alk in fly

#### Fly husbandry and phenotyping

For fly crosses, 15 virgin females were crossed with 3 males in bottles containing 1.55% live yeast, cornmeal, sugar, and agar.^52^ Crosses were dumped 5 days following crossing, and female progeny were sorted into 4 replicate vials of 25 flies each, with food containing 200μm RU486 to induce activation of the Gal-UAS system.^53^ Flies were maintained in 65% relative humidity at 25°C in a 24-hour light/dark cycle throughout life. Two weeks post-induction, phototaxis was tested as previously described^38^ by placing flies in a clear, empty 30 cm.-long vial horizontally in a dark room. Light was shined on one end and the number of flies in the last 10 cm. closest to the light source after 1 minute was scored for responsiveness to light signals. This was tested across each of the 4 vials per group in 3 biological replicates (total 100 flies per replicate). Strains used were 3x*elav*-GS (provided from the lab of Geetanjali Chawla)^54^ for RU486-dependent pan-neuronal Gal4, *w*^*Dah*^ control strain, UAS-*Alk*^RNAi^, and UAS-*Alk*^DN^ (provided from the lab of Linda Partridge)^23^.

## Acknowledgments

We would like to thank Ajay Kumar for critically reviewing our manuscript. This research has been conducted with the UK Biobank resource application 17643. KAW is supported by NIH T32AG000266-23. We thank the Bloomington Drosophila Stock Center for providing flies used in this study. This work is funded by grants awarded to P.K. from the Reta Haynes Foundation, American Federation of Aging Research, NIH grants R01 R01AG038688 and AG045835 and the Larry L. Hillblom Foundation.

## Author Contributions

S.A., M.B., A.B., A.V. and P.K. designed research; K.A.W., E.C. performed experimental validations;

S.A., K.A.W., B.B., A.V., C.Y.M., D.B., O.P., M.B., A.B. and P.K. analyzed data;

S.A., K.A.W., B.B., A.V., C.Y.M., R.L., J.S., M.B., A.B. and P.K. contributed to the interpretation of the results;

S.A., K.A.W., B.B., A.V., C.Y.M., A.B. and P.K. wrote the paper.

