## Supplemental Materials for "Longitudinal fundus imaging and its genome-wide association analysis provide evidence for a human retinal aging clock"

### Supplemental Information

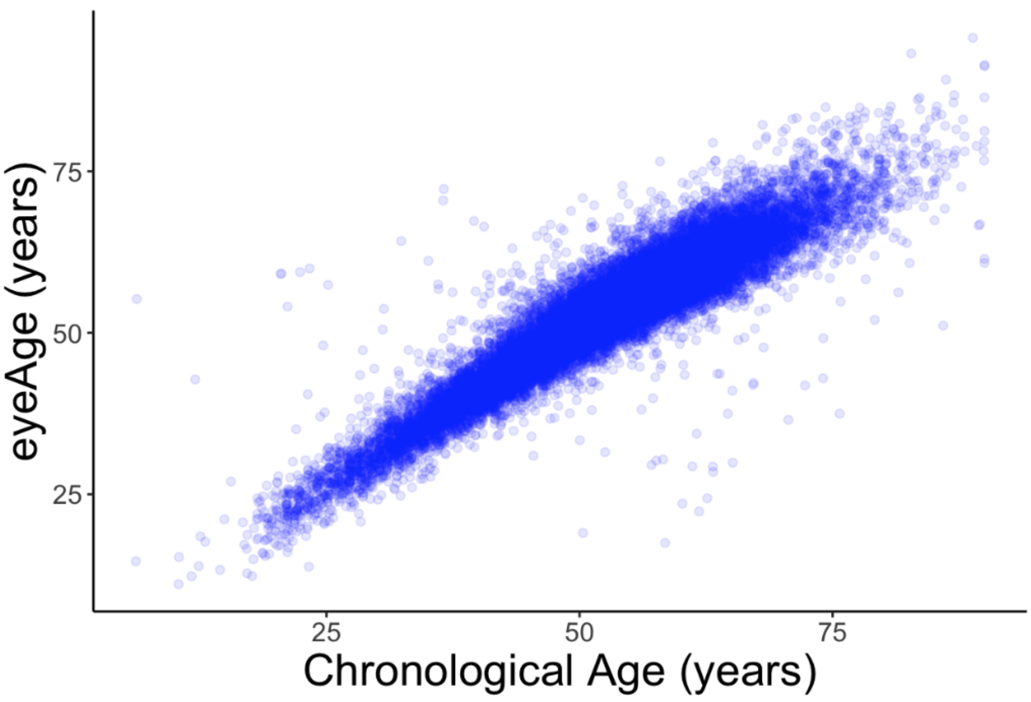


**Supp Figure 1. Scatter plot of eyeAge with chronological age (Pearson $\rho$ = 0.96)**

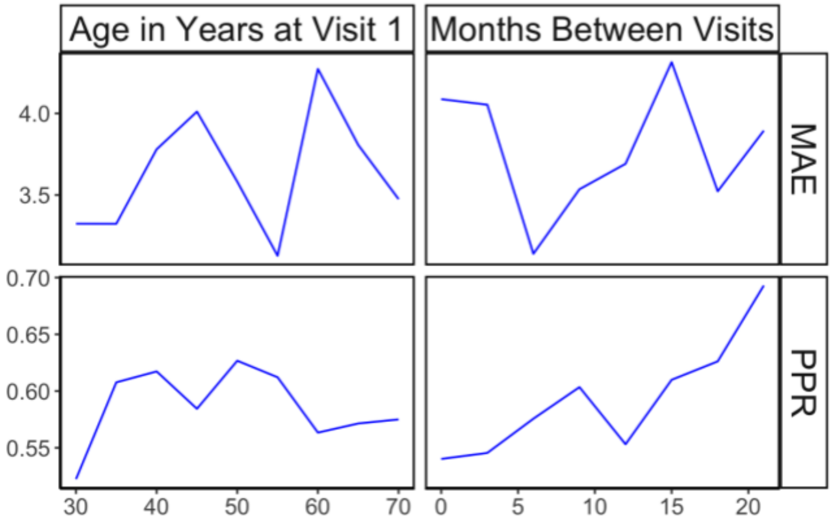


**Supp Figure 2. Positive prediction ratio and MAE for random, time-matched individuals.** Plots shown in relationship to chronological age (left) and time between longitudinal visits (right).

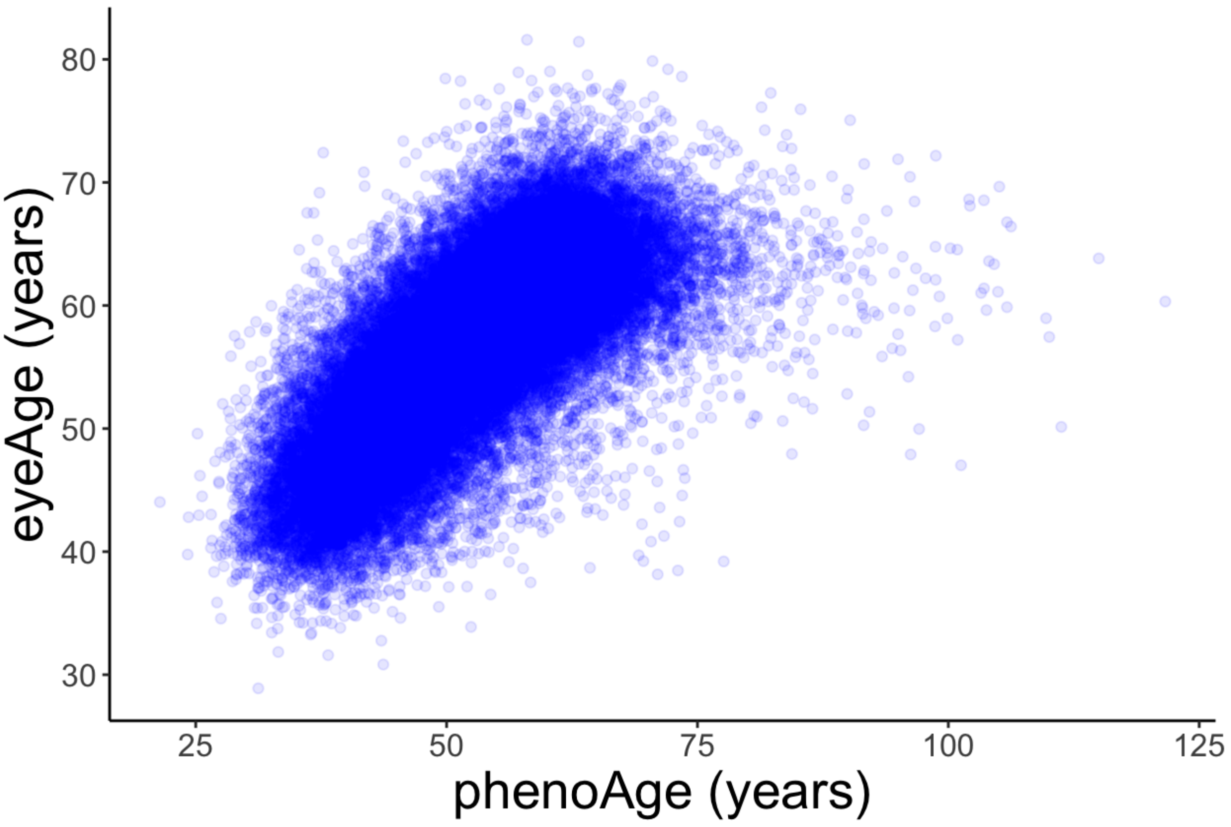


**Supp Figure 3. Scatter plot of eyeAge and phenoAge (Pearson $\rho$ = 0.71)**

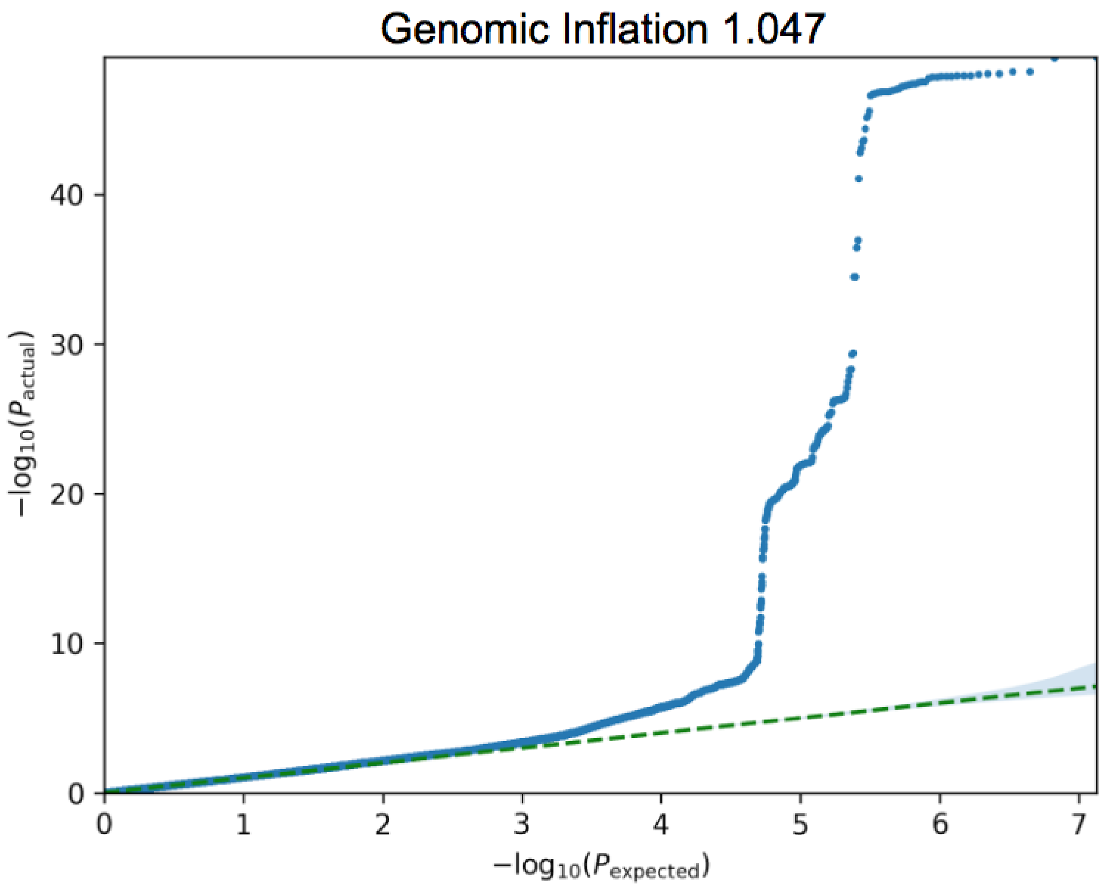


**Supp Figure 4. eyeAgeAcceleration qq-plot.**

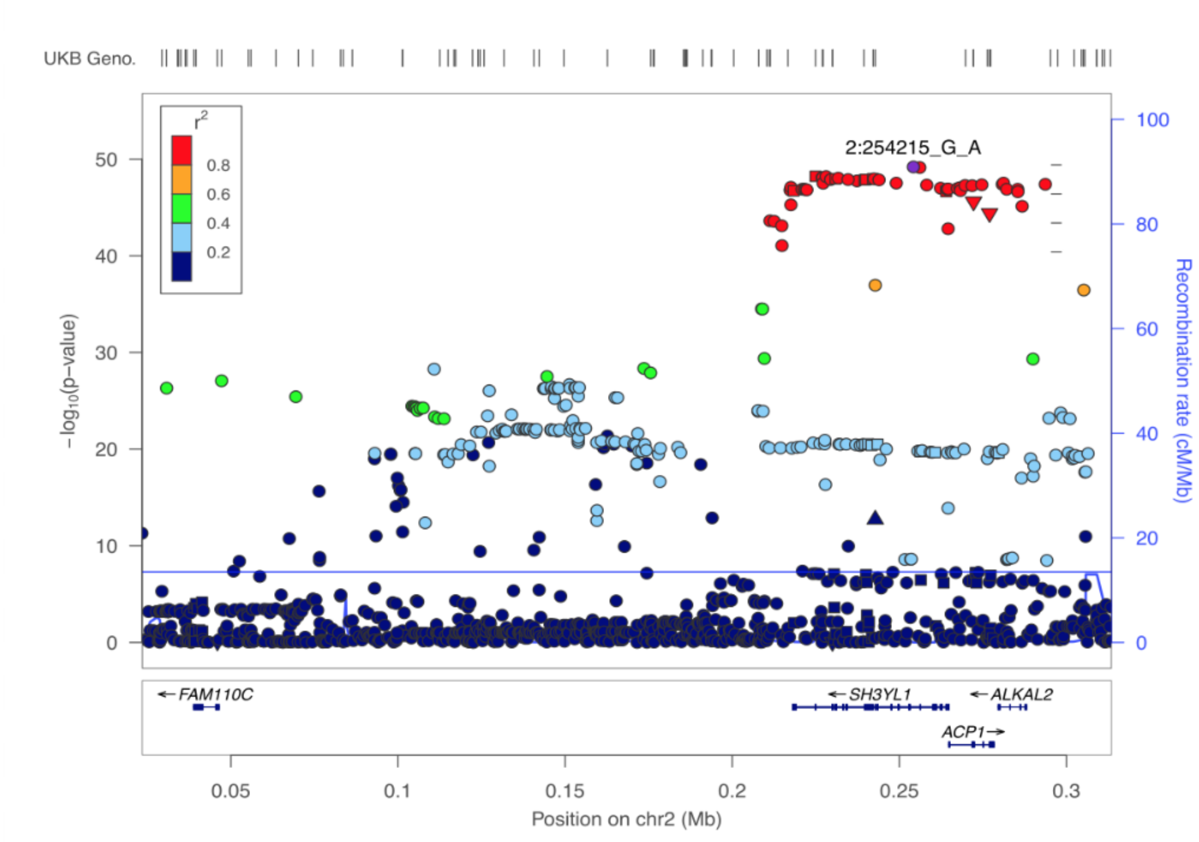


**Supp Figure 5. Zoom in on significant locus covering three genes in a highly significant LD block**. This block includes the three genes: *SH3YL1*, *ACP1*, and *ALKAL2.*

### Supplemental Tables

Supplemental Table 1: Association results_filtered
Supplemental Table 2: Fine mapping
Supplemental Table 3: Association results_annotated_hits
Supplemental Table 4: genes associated with eyeAgeAccel and function
