## Supplemental Table 4 for "Longitudinal fundus imaging and its genome-wide association analysis provide evidence for a human retinal aging clock"

**Supplemental Table 4.** List of genes associated with eyeAgeAccel and function

| Gene | P value | Function/disorder association | Reference |
|---|---|---|---|
| SH3YL1 | 6.40E-50 | Phosphatidylcholine binding; type 2 diabetes | [1] |
| OCA2 | 1.90E-12 | Eye pigmentation | [2] |
| POC5 | 6.90E-10 | Centrosome interactions; macular degeneration | [3] |
| GJA3 | 1.60E-09 | Gap junction protein; cataracts | [4] |
| MEF2C | 1.70E-09 | Myocyte enhancement; Alzheimer's disease, muscle wasting | [5,6] |
| TSPAN11 | 2.20E-09 | Cell migration; cancer prognosis | [7] |
| NKX6-1 | 5.80E-09 | Beta cell development; cancer prognosis | [8] |
| GRM5 | 1.30E-08 | Metabatropic glutamate receptor; hearing loss | [9] |
| SLC16A1 | 2.20E-08 | Monocarboxylic acid transporter; cancer prognosis | [10] |
| GPR139 | 3.30E-08 | Rhodopsin family GPCR; brain metabolism with age | [11] |
| ARMS2 | 4.00E-08 | Eye extracellular matrix activity; macular degeneration | [12] |

| | | | |
|---|---|---|---|
| RAET1G | 7.70E-08 | Retinoic acid activity; cancer prognosis | [13] |
| ACP1 | 1.20E-07 | Phosphotyrosine protein phosphatase activity; retinopathy, diabetes, lifespan | [14,15] |
| ALKAL2 | 2.80E-07 | Tyrosine kinase activity; lifespan | [16] |
| SNTG1 | 3.00E-07 | Gamma enolase trafficking activity; Alzheimer's disease, stroke, cancer prognosis | [17-19] |
| BAZ2B | 3.30E-07 | Chromatin remodeling; Alzheimer's disease | [20] |
| GJD2 | 3.40E-07 | Gap junction activity; myopia | [21] |
| JAG1 | 3.70E-07 | Notch ligand activity; retinopathy, coronary artery disease | [22,23] |
| MYT1 | 5.20E-07 | Myelin transcription factor; age-related neuron demyelination | [24] |
| RASGRF1 | 5.70E-07 | Guanosine nucleotide exchange factor; myopia, neurogenesis | [25,26] |
| ARRDC3 | 6.20E-07 | Arrestin-related GCPR activity; Alzheimer's disease, glioma, breast cancer | [27-29] |
| RASSF3 | 6.60E-07 | GTP binding activity; glaucoma | [30] |

| | | | |
|---|---|---|---|
| COL4A3 | 7.10E-07 | Collagen activity; Alport syndrome, macular degeneration | [31,32] |
| DIRC3 | 8.50E-07 | Tumor suppressor activity; carcinoma prognosis | [33] |
| LZTFL1 | 8.80E-07 | Transcription factor activity; retinal degeneration | [34] |
| ATP6V0A2 | 9.30E-07 | ATPase activity; cutis laxa | [35] |
| GCNT3 | 9.80E-07 | Glycosyltransferase enzymatic activity; cancer prognosis | [36] |
| COL4A4 | 1.00E-06 | Collagen activity; Alport syndrome | [52] |
